# Glucosinolates promote initial population establishment of feral oilseed rape

**DOI:** 10.1101/429290

**Authors:** Elze Hesse, Dave J. Hodgson, Tom J. de Jong

**Affiliations:** Institute of Biology, University of Leiden, PO Box 9505, 2300 RA Leiden, The Netherlands; Biosciences, University of Exeter, Penryn Campus, Penryn, TR10 9FE, UK

**Keywords:** Competition, Crop ferality, Glucosinolates, Herbivory, Population dynamics

## Abstract

**Background:** Crops are often selected for traits that confer a selective disadvantage in the wild. A key trait that has been greatly altered by domestication is investment in herbivore defence. It remains unclear, however, whether variation in chemical defence affects a crop’s ability to colonize semi-natural habitats where it typically has to compete with a resident community. Here, we investigate how breeding efforts aimed at reducing glucosinolate levels in seeds – canonical herbivore deterrents – influence initial establishment of *Brassica* populations spanning a wild-feral-domesticated gradient.

**Methods:** We followed the dynamics of twenty-nine *Brassica* accessions in two experimental fields by recording life table parameters and vegetation cover biannually over a two-year period. Accessions were selected to vary in their glucosinolate content, and included lines of wild turnip (*B. rapa)*, feral *B. napus* as well as modern canola and historical oilseed rape cultivars. Populations were established by sowing seeds on bare soil after which the natural vegetation was allowed to regenerate, providing a temporal gradient in the degree of interspecific competition.

**Results:** Populations flourished in the first year, but many perished during a second year of growth, in particular those of oilseed rape. Declines coincided with an increase in vegetation cover, but were slower in populations harbouring more glucosinolates. These compounds had opposing effects on different life cycle stages: seedling establishment was greater in high-glucosinolate lines, which traded off with reduced post-recruitment survival. Crucially, the effect of glucosinolates on persistence was lost when focussing on oilseed rape only, but the underlying demographic trade-off remained.

**Discussion:** Our study illustrates that initial establishment of feral oilseed rape is governed by glucosinolate-mediated trade-offs between seedling recruitment and subsequent survival, with low-glucosinolate lines (modern canola) being most successful when post-recruitment conditions are relatively benign. Such demographic trade-offs likely extend to other species, and must be considered when managing escaped crops and invasive plants.

## Introduction

Crops are selected for traits to improve human consumption and yield in agricultural settings (Gepts 2004; McKey et al. 2012). Selection for human-beneficial traits is predicted to hamper crop ferality and establishment in more natural habitats (Warwick & Stewart 2005). A key trait that has been greatly altered by domestication is investment in secondary metabolites; these chemical compounds are often selected against because of their anti-nutritional value and/or toxicity (Meyer, DuVal & Jensen 2012; Chen et al. 2015). The presence of these compounds deters many generalist herbivores (Futuyma & Agrawal 2009); hence crops are therefore likely more susceptible to herbivore attack compared to their wild ancestors (Turcotte, Turley & Johnson 2014; Simpson et al. 2017). Crucially, selection against herbivore deterrents might come at a cost of reducing population persistence when herbivores attack key life stages (Strauss et al. 2009). Here, we test this outstanding issue by comparing the demography of *Brassica* lines varying widely in their domestication history and glucosinolate profiles (Hopkins, van Dam & van Loon 2009).

*Brassica napus*(oilseed rape) is widely grown in temperate regions across the globe. Feral oilseed rape is pervasive, forming transient populations that persist via repeated influx of seeds (Pivard et al. 2008; Devos et al. 2012; Meffin, Duncan & Hulme 2015). Congruent with other crops (Wang et al. 1999; Gepts 2004; Gressel 2008; Meyer, DuVal & Jensen 2012; Meyer & Purugganan 2013; Milla et al. 2015), oilseed rape has become increasingly differentiated from its wild ancestors *Brassica rapa* and *B. oleracea*(Gepts 2004). In particular, breeding programs have selected against anti-nutrients in oil, such that seeds of modern canola lines typically contain ≤ 2% erucic acid (EA, % total oil) and ≤ 30 µmoles of glucosinolates (GS, g^-1^ air-dried oil-free meal) (Olsen & Sorensen 1980). Crucially, GS play a key role in herbivore deterrence (Hopkins, van Dam & van Loon 2009) and recent evidence pinpoints to a minor defensive role of EA against seed predators (de Jong et al. 2016). Understanding how variation in chemical defence affects oilseed rape ferality requires identification of ecological conditions that permit initial establishment from seeds. While theoretical models have aimed at addressing this outstanding issue (Claessen et al. 2005; Garnier & Lecomte 2006), demographic data encompassing the entire life cycle are scarce (Pessel et al. 2001; Bagavathiannan & Van Acker 2008; Pascher et al. 2010; but see: Crawley et al. 1993; Crawley & Brown 1995; Crawley & Brown 2004; Hooftman et al. 2015). This is an important knowledge gap as oilseed rape is a major crop for which transgenic lines have been developed (Ellstrand, Prentice & Hancock 1999; Hails & Morley 2005; Warwick et al. 2008; Schafer et al. 2011; Bailleul, Ollier & Lecomte 2016).

Defence compounds are thought to play a key role in the invasion success of wild Brassicas (Muller 2009) and there is some evidence this also holds for oilseed rape: slugs greatly prefer modern canola seedlings over more historical high-GS lines (Glen, Jones & Fieldsend 1990; Giamoustaris & Mithen 1995; Moshgani, Kolvoort & de Jong 2014). Because recruitment is a crucial point in the life cycle of many annual herbs, herbivore deterrence at this stage could potentially increase persistence (Strauss et al. 2009). However, whether seedling mortality comes at a cost of reducing persistence remains unclear as many ruderal plants rely on disturbance to sustain viable populations by providing bare sites for seedling recruitment (Grime 1977; Crawley & Brown 1995; Levine, Adler & Yelenik 2003; Crawley & Brown 2004; Duncan et al. 2014).

Here, we investigate how variation in GS and competition jointly affect establishment of *Brassica* by collecting demographic data in 29 experimental populations spanning a wild-feral-domesticated gradient, including lines of *B. rapa*, feral *B. napus* as well as modern canola and historical oilseed rape cultivars. Owing to their low GS content, we expect canola seedlings to experience relatively high levels of herbivory, which should reduce population viability in habitats where recruitment contributes significantly towards persistence. Conversely, we expect historical lines to be more akin to *B. rapa* in terms of GS, herbivory and persistence. Little is known about selection on herbivore defence in feral oilseed rape. Agricultural settings often present strong ecological contrasts with more natural habitats (Denison, Kiers & West 2003; Chen, Gols & Benrey 2015): while these habitats share many natural enemies (Blitzer et al. 2012), herbivores likely pose weaker selection on defensive traits in agricultural systems, where humans typically modulate their presence (e.g. molluscicide application to reduce seedling mortality: Garthwaite et al. 2013). However, the relatively short life span of many feral populations (but see Charters, Robertson & Squire 1999; Pessel et al. 2001; Pascher et al. 2010) in combination with reduced genetic diversity in crop more generally (Doebley, Gaut & Smith 2006) likely impedes selection on defence traits. Hence, canola and feral oilseed rape likely display similar GS profiles, herbivory and persistence.

## Material and Methods

### Study populations

Twenty-nine *Brassica* lines have been selected to vary in their EA and GS content, including modern canola (C = ‘Hornet’, ‘Oase’, ‘Lioness’, and ‘Billy’ were obtained from the plant breeding company DSV, ‘Ladoga’ from Limagrain, and two ‘Pioneer’ cultivars from DuPont) and historical oilseed rape cultivars (CO = ‘Windal’, ‘Velox’ and ‘Mansholt’s Hamburger’ date from 1959, 1967 and 1899, respectively, and were obtained from the Centre for Genetic Resources in Wageningen). In summer 2009, we also collected seeds in populations of feral oilseed rape (BN; *n* = 10 populations) and *B. rapa* at different localities in the Netherlands (BR; *n* = 9 populations). Seeds were combined into a single-population bulk sample (sample sizes and localities provided in Table S1).

For each accession, GS and EA were extracted from ten seeds. Aliphatic GS were identified using HPLC spectrometry with sinigrin as an external standard (van Dam, Witjes & Svatos 2004). GS are present in other plant tissues, albeit at lower concentrations (Stefansson & Downey 1995): aliphatic GS content of seeds and leaves was strongly correlated in our study lines (Spearman’s correlation, ρ = 0.83, *P*< 0.001), with leaves of modern canola typically containing < 4 µmol g^-1^ dry weight. The content of EA in oil was analysed using the methods described in Rücker and Röbbelen (1996). In line with oilseed rape’s putative breeding history, we found that seeds of historical cultivars typically contained high EA and aliphatic GS levels, comparable to those observed in *B. rapa*(Fig. 1). As expected, modern canola varieties generally had relatively low and invariable EA and aliphatic GS contents. Five feral populations displayed higher GS levels than any of the modern canola varieties tested (Fig. 1). Indole GS content did not vary across accessions (data not shown).

**Figure 1.**
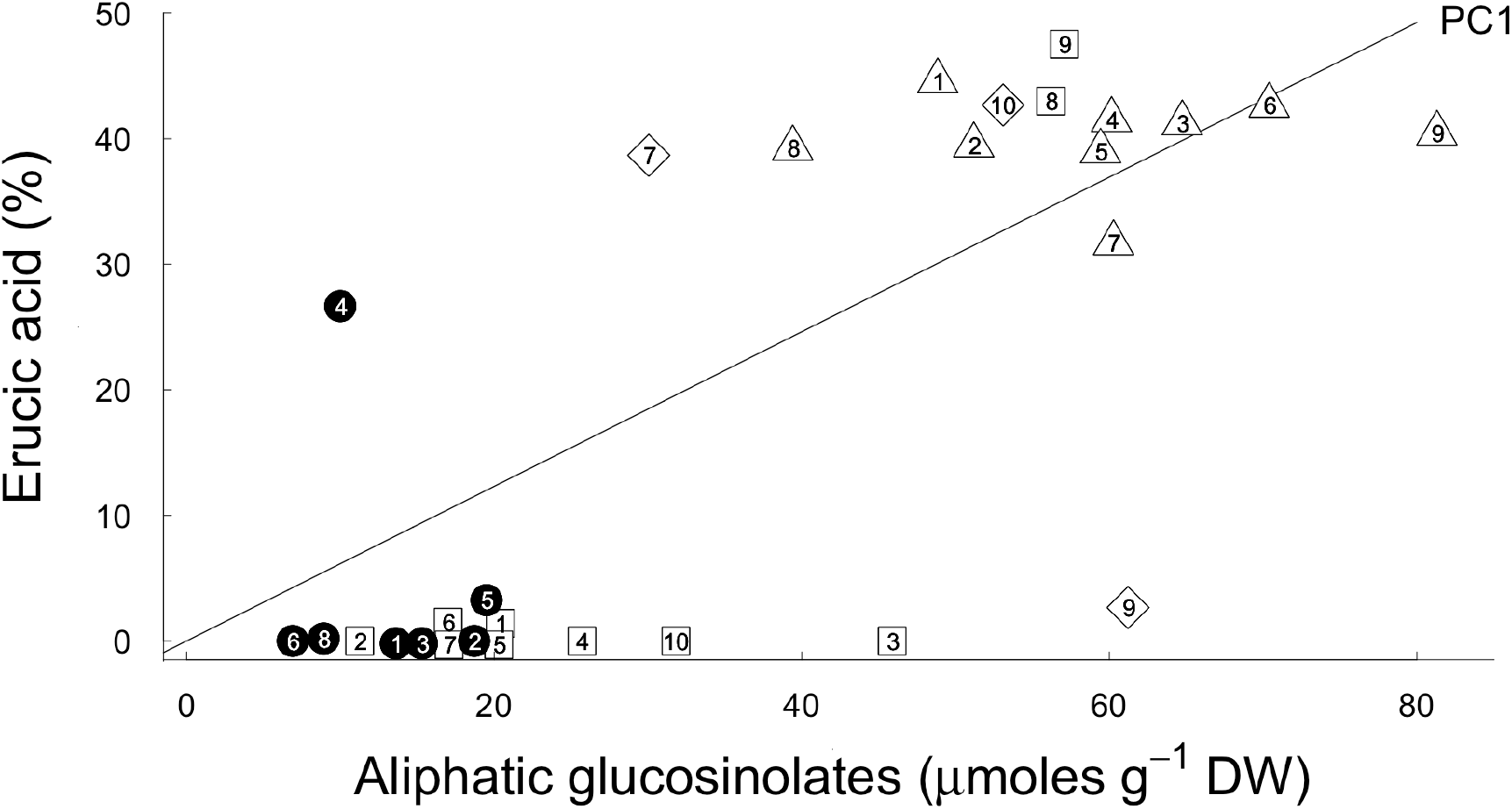
Relationship between mean erucic acid (EA, % of seed oil) and aliphatic glucosinolate (GS, µmol g^-1^1 air-dried oil-free meal) content of seeds, where dots = C, diamonds = CO, squares = BN and triangles = BR. The fitted line depicts the first principal component (‘chemical PC1’) derived from a PCA analysis, accounting for 95% of original variance. More detailed information about the study populations can be found in Table S1.

### Data collection

Populations were established in late August 2009 by sowing seeds on tilled soil in two experimental fields in the Netherlands (Boskoop: 52°05’04.5” N, 4°41’20.1” E and Swifterbant: 52°31’28.4”N, 5°38’11.0”E). The local vegetation comprised tall herb-and grass communities and was allowed to regenerate over the course of our study. Study species were naturally absent in our experimental fields, but populations could be found in close proximity along river-and roadsides. In order to minimize potential cross-pollination, lines were segregated into blocks that were separated from one another by a buffer zone of 4 meters. In each block, 100 seeds of a single accession were sown in eight replicate 1-m^2^ plots. To account for block effects, a control plot was sown with 100 seeds of the *B. napus* cultivar ‘Nootzoet’ (Vreeken’s Zaden); however, no such effects were apparent (data not shown). We observed that bees mainly visited flowers *within* plots before moving on to other patches, suggesting that gene flow between populations was restricted, which was confirmed by flow cytometry: out of 130 second-cohort plants analysed for chromosome numbers, only three plants were hybrids.

Within each plot, life table parameters were measured biannually over a two-year period (Autumn 2009–2011), including the number, identity, and size of plants and their state (alive/dead, reproductive/non-reproductive) as well as bare soil cover. Because populations attained particularly high densities after self-seeding, we marked a subset of plants for the next cohort, and biannually recorded the stage-specific number of individuals. In early spring, we also scored the degree of herbivore damage, ranging from no damage (0) to almost completely consumed (3). For each accession, fecundity was quantified for a subset of plants, using estimates of seed set, fruit numbers and pre-dispersal seed predation (Table S1). Because the majority of predated fruits did not produce any viable seeds, pre-dispersal predation was estimated as the proportion of fruits displaying any sign of insect-induced damage. Seedling establishment was calculated as the number of seedlings emerging in autumn/spring divided by the number of seeds entering a plot (via sowing or self-seeding).

### *The life cycle of* Brassica

The two *Brassica* species largely behave as winter annuals with relatively simple life cycles: seedlings establish in autumn or in the following spring, flowering and seed ripening occur during summer and seed is shed in early autumn. Our assumptions on the demographic processes taking place over the course of our study are summarised in Fig. S1. As for most crops, seed bank survival is limited compared to that of wild relatives (Hails et al. 1997), yet a small fraction of oilseed rape seeds can remain dormant for up to ten years (d’Hertefeldt, Jørgensen & Pettersson 2008). Seed dormancy has often been invoked in playing a crucial role in driving oilseed rape ferality (Devos et al. 2012; Hooftman et al. 2015), yet the build-up of a seed bank is limited without soil disturbance (de Jong, Tudela Isanta & Hesse 2013). The majority of seeds germinate immediately upon reaching the soil surface if given adequate moisture (Lutman et al. 2005), thereby preventing the development of dark dormancy, and hence the incorporation of seeds into a soil seed bank. Because plots were left undisturbed over the course of our study, we assume a negligible contribution of seed bank recruitment to population growth. This was verified by collecting topsoil (∼5 cm) from all demographic plots at the end of our study: very few seedlings of either species emerged from the soil after a 14-day germination period under optimal conditions (*B. rapa*= 10 and 3; *B. napus* = 3 and 1 in Boskoop and Swifterbant, respectively).

The finite rate of increase (*λ*) can therefore be calculated as: *λ* =*rs_a_F,* where vegetative plants have a probability *s_a_* to reach the flowering stage, producing a mean number of *F* seeds, and *r* is cumulative seedling recruitment, which can be decomposed into *r = r_a_s_w_ + r_s_*. The first term is the fraction of newly emerged seedlings in autumn *r_a_*, which survive over winter as non-reproducing individuals *s_w_*, and the second term is spring seedling emergence *r_s_*. Note that *B. napus* seeds readily germinate upon reaching the soil surface, with germination fractions being close to 100%. Because germinants might have died before being recorded, our measure captures successful recruitment rather than germination rates.

### Data analyses

All analyses were implemented in R (v. 2.2.2 R Development Core Team). Because many second-cohort Boskoop populations perished during the early life stages (Table S1), *λ* was set to a minimum threshold of 0.001 for those populations in all subsequent analyses. We first tested how competition and breeding history (C = modern canola, CO = old cultivars, BN = feral oilseed rape and BR = *B. rapa*) influenced *λ*, using a general linear model (GLM) with bare soil cover, breeding history, site and year as explanatory variables (model 1). The full model considered the three-way interaction between history, site and year as well as all twoway interactions. To obtain a proxy for annual vegetation turnover, we carried out a principal components analysis (PCA) on spring and autumn estimates of bare soil; the first principal component (‘soil PC1’) explained 90% of the original variance and was used in all analyses as a covariate. Using a similar approach (model 2), we next tested whether investment in chemical defence plays a key role in driving persistence by substituting ‘history’ with mean population-level EA/GS content, which was summarized by the first principal component of a PCA on these chemical traits (‘chemical PC1’; Fig. 1). The most parsimonious model was arrived at by sequentially deleting terms and comparing model fits using F-tests, after which Tukey-adjusted least-squares means, slopes and their contrasts were computed using the ‘lsmeans’ package (Lenth 2016).

To disentangle whether variation in persistence is driven by domestication *per se* or by chemical defence in particular, we used a linear mixed effects model (model 3) with ‘soil PC1’, site, year and ‘chemical PC1’ as fixed effects and random intercepts fitted to different levels of breeding history (‘lmer’ function in lme4 package; Bates et al. 2015). The full model considered the three-way interaction between site, year and ‘chemical PC1’ and all two-way interactions. This allowed us to test for the effect of chemical PC1 while accounting for other systematic differences associated with domestication. Full models were simplified by sequentially eliminating non-significant terms and interactions (P > 0.05) following a stepwise deletion procedure. The significance of the explanatory variables was established using Wald-type likelihood ratio tests, which were *χ*^2^ distributed. We used a similar tiered approach to disentangle the effects of domestication, competition and secondary metabolites on life stages. To achieve normality of standardized residuals and equal homogeneity of variance, variables were transformed where appropriate (see results). To test whether patterns were mainly driven by cross-species differences we also carried out analyses using the oilseed rape data set only.

To identify demographic rates that contribute significantly towards variation in *λ*, we carried out a modified key factor analysis by regressing yearly vital rates *k_i_* on *λ*: *log(k_i_)* = *β*_*o*_ + *β*_1_*log(λ)*. The linear regression coefficient, *β*_*1*_, indicates to what extent differential performance in life stage *i* leads to corresponding changes in *λ* (Podoler & Rogers 1975); the *β*_*1*_’s for all vital rates sum up to one and rates with the largest values of *β*_*1*_ are by definition considered key factors. To avoid the logarithm of zero, one individual was added to vital rate calculations where appropriate. While mortality-based key factor analyses have certain drawbacks (reviewed in Royama 1996), their conceptual problems can be overcome by assessing the effect of life-history variation on *λ* (Sibly & Smith 1998).

## Results

*The role of domestication and chemical traits on life cycle transitions and their contribution towards* Brassica *persistence*

Bare soil cover decreased over time and varied between experimental fields: plots in Swifterbant were less densely vegetated compared to those in Boskoop (Fig. 2A) throughout the course of our study. All populations thrived in the first year, but λ decreased dramatically during a second year of growth (Fig. 3A), which coincided with a reduction in bare soil cover (soil PC1 effect in models 1–3: F_1, 103_= 5.68, P < 0.05; F_1, 111_= 37.83, P < 0.001 and *χ*^2^_1_= 41.56, P < 0.001; Fig. 2A and Table S2). The rate of population decline varied as a function of breeding history, being exceedingly faster in cultivated oilseed rape compared to feral *B. napus* and/or wild *B. rapa* (history x year in model 1: F_3, 103_ = 10.39, P < 0.001; Fig. 3A). Accessions performed differentially depending on their growing locality (history x site in model 1: F_3, 103_ = 5.30, P < 0.01): *B. rapa* populations outperformed those of oilseed rape, but only in Boskoop where cultivated *B. napus* attained overall lower densities (Table S2). Importantly, yearly changes in *λ* were not only driven by competition, but also by variation in chemical defence (year x chemical PC1 in model 2: F_1, 111_ = 13.38, P < 0.001; Fig. 3A and Table S2). While EA and GS did not affect *λ* of the first cohort, populations with greater investment in these compounds outperformed those investing relatively little during a second year of growth (Fig. 3A). When accounting for systematic differences associated domestication, which explained ∼25% of the observed variation, these effects were still apparent (year x chemical PC1 in model 3: *χ*^2^_1_ = 15.40, P < 0.001; Table S2).

**Figure 2.**
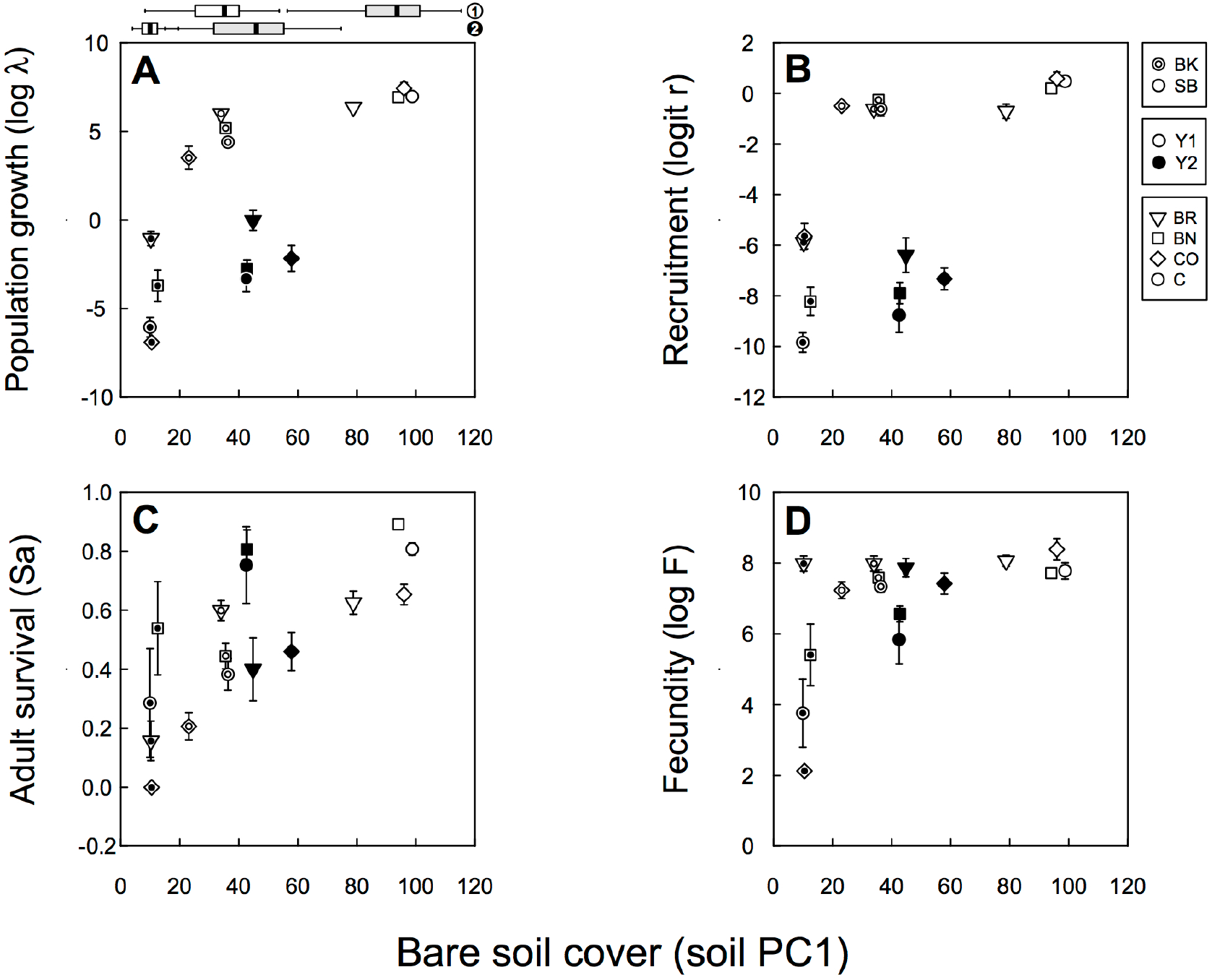
Plotted relationship between mean ± SE bare soil cover (soil PC1) and demographic transitions for each accession type (dots = C, diamonds = CO, squares = BN and triangles = BR), where (A) population growth, (B) seedling recruitment, (C) survival to flowering and (D) fecundity during the first (open symbols) and second (closed symbols) year of our study in Boskoop (BK) and Swifterbant (SB). Temporal changes in bare soil cover (1 = 1^st^ year and 2 = 2^nd^ year) in Boskoop (open boxplots) and Swifterbant (grey boxplots) are provided on top of panel A.

**Figure 3.**
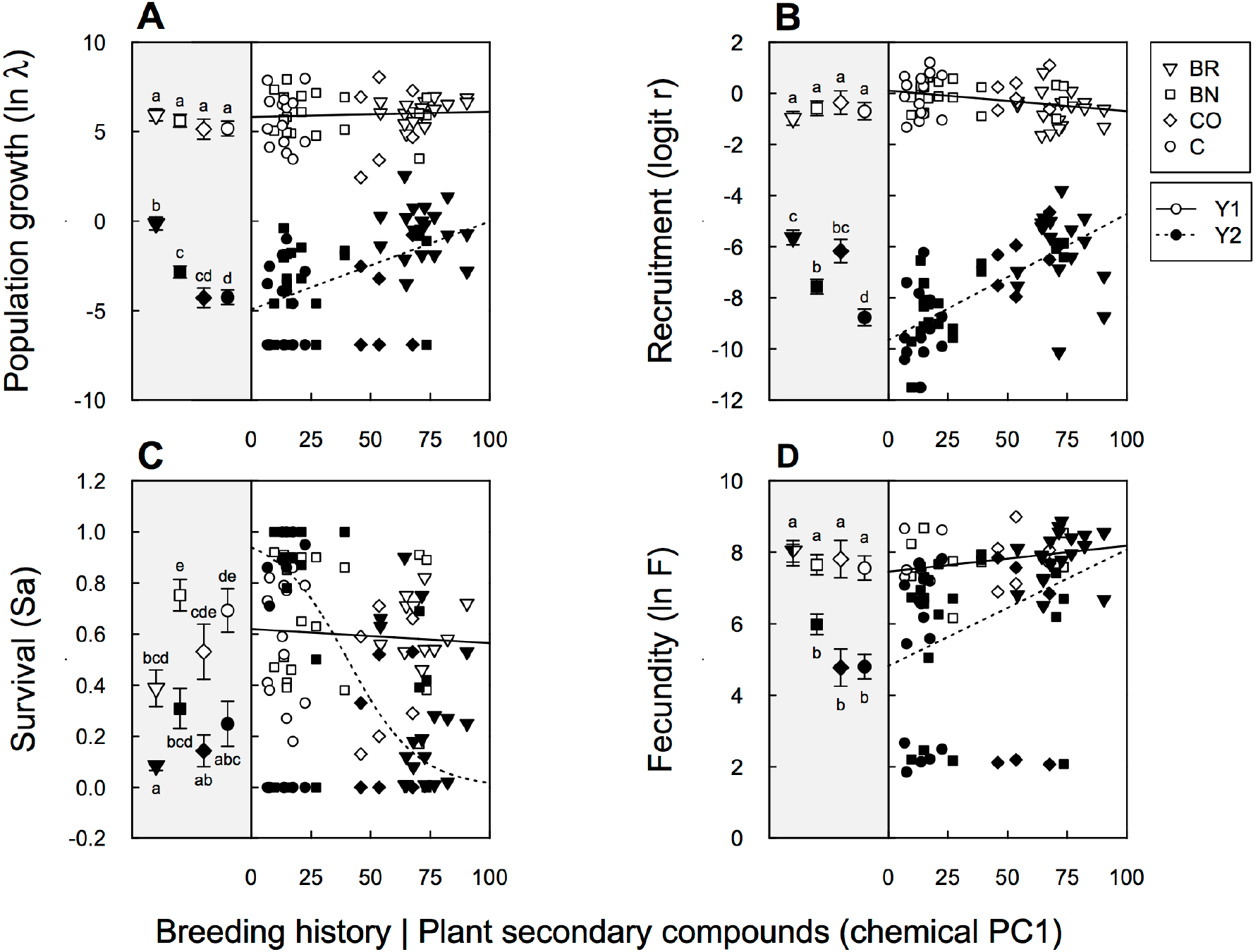
The influence of breeding history and plant secondary compounds (chemical PC1) on demography, where: (A) population growth, (B) seedling recruitment, (C) fraction of established plants surviving to the flowering stage and (D) fecundity in the first (open symbols, solid line) and second year (solid symbols, dashed line) of our study. Letters denote significant differences between pairwise Tukey-adjusted least-squares estimates (dots = C, diamonds = CO, squares = BN and triangles = BR; model 1). Lines depict temporal changes in the relationship between chemical PC1 and demographic transitions, the parameter estimates of which are given in Table S2 (model 2).

Recruitment patterns mirrored those found for *λ* (Fig. 3B). Seedling recruitment was negatively affected by a reduction in bare soil cover (soil PC1 in models 1–3: F_1, 106_= 13.49, P < 0.001; F_1, 109_= 13.41, P < 0.001 and *χ*^2^_1_= 13.50, P < 0.001; Fig. 2B and Table S2). In addition, declines in recruitment were dependent on breeding history, being strongest in modern canola, followed by feral oilseed rape/historical cultivars and finally *B. rapa* (history x year in model 1: F3, 106 = 13.64, P < 0.001; Fig. 3B and Table S2). The presence of EA and GS increased seedling establishment (Fig. 3B), the extent of which varied across experimental fields (site x chemical PC1 in model 2: F1, 109 = 8.25, P < 0.01) and years (year x chemical PC1 in model 2: F1, 109 = 59.86, P < 0.001), being greater in Boskoop and in the second year of our study (Table S2). The relationships between EA/GS and recruitment remained significant after accounting for other systematic differences arising from domestication (model 3: chemical PC1 interaction with year and site, respectively: *χ*^2^_1_ = 50.77, P < 0.001 and *χ*^2^_1_ = 13.50, P< 0.01; Table S2).

Plant survival differed between sites (models 1 and 2 with quasibinomial error structure: F1, 101 = 5.71, P < 0.01 and F1, 105 = 21.46, P < 0.001, respectively), being significantly lower in Boskoop, irrespective of bare soil cover (P > 0.05 in models 1–3; Fig. 2C and Table S2). Temporal changes in survivorship were related to breeding history (year x history in model 1: F3, 101 = 5.71, P < 0.01): survival of oilseed rape lines was reduced to a greater extent compared to that of *B. rapa* during a second year of growth (Fig. 3C). Importantly, survival was significantly lower in lines harbouring more EA/GS, in particular in the second year of our study (year x chemical PC1 in model 2: F1, 105 = 21.82, P < 0.001). The negative effect of EA/GS on survivorship persisted when accounting for breeding history and was strongest for the second cohort of plants growing in Boskoop (model 3: site x year x chemical PC1: *χ*^2^_1_ = 5.30, P < 0.05; Table S2).

Seed production was largely independent of vegetation cover (soil PC1 effect in models 1–3: F1, 99 = 2.68, F1, 108 = 1.48, *χ*^2^_1_= 2.91, all P > 0.05; Fig 2D). Seed production differed as a function of breeding history, the extent of which varied between years (history x year model 1: F3, 103 = 7.38, P < 0.001; Table S2) and sites (history x site model 1: F3, 103 = 5.21, P < 0.01; Table S2). While first-cohort plants produced similar seed numbers, oilseed rape fecundity was reduced to a great extent in the second year of our study, in particular that of cultivars growing in Boskoop (Fig. 3D and Table S2). There was temporal variation in seed production across sites (site x year in models 1–3: F1, 103 = 5.56, P < 0.05; F1, 110 = 4.14, P < 0.05 and *χ*^2^_1_ = 4.59, P < 0.05), with fecundity being lowest for the second generation of plants growing in Boskoop (Table S2). Crucially, fecundity was greater in lines harbouring more EA/GS, but only so in the second year of our study (year x chemical PC1 in models 2 and 3: F1, 110 = 6.45, P < 0.05 and *χ*^2^_1_ = 7.08, P < 0.02; Fig. 3D and Table S2). Below we discuss key differences between cross-species and oilseed rape only analyses; a summary of the differential effects of GS on various life cycle transitions is depicted in Fig. 4.

**Figure 4.**
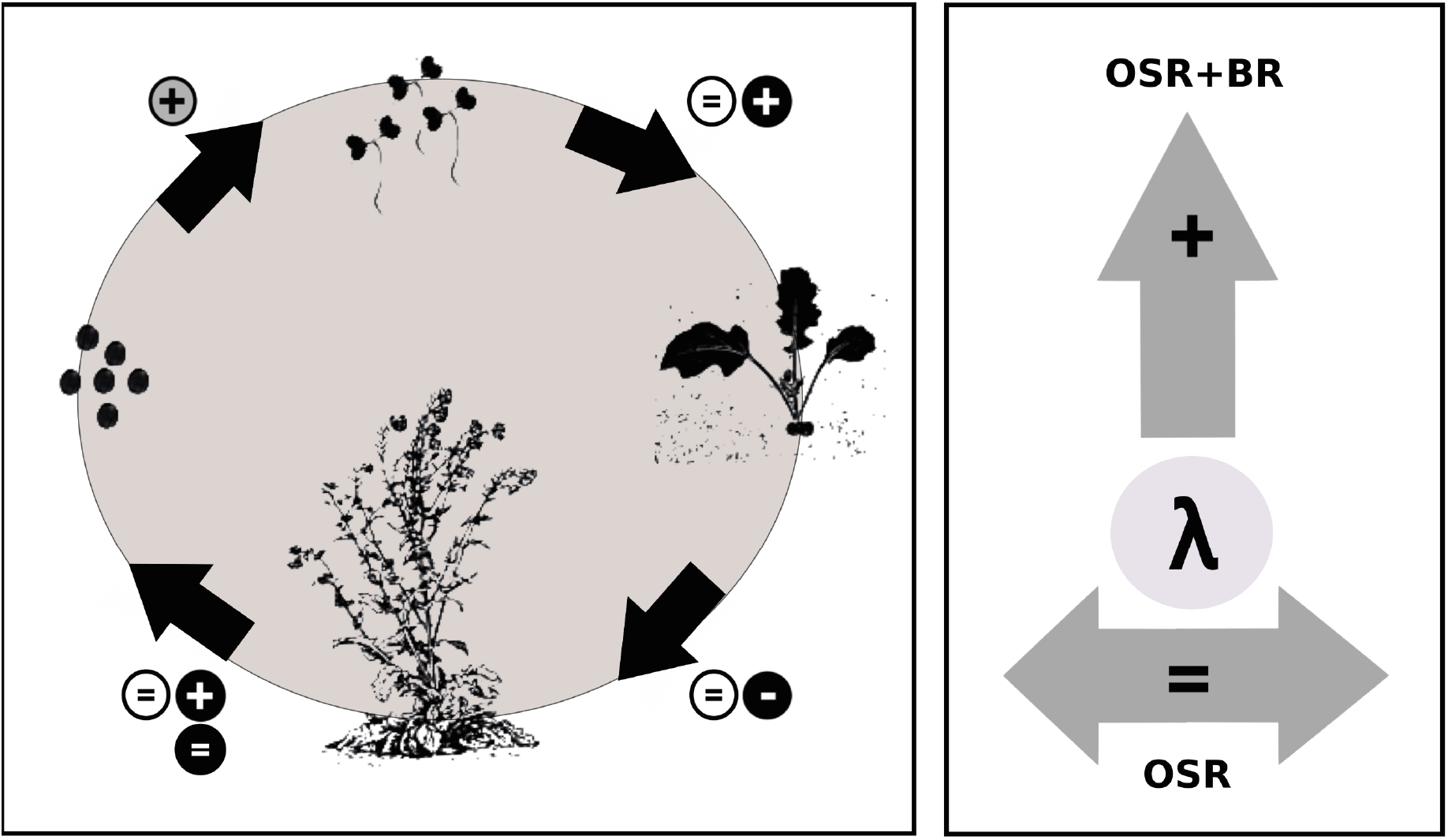
Illustrative graph summarizing the effect of glucosinolates on different life cycle transitions in the first (open circles) and second year (filled circles) of our study and their net effect on population growth (λ) based on the cross-species (OSR+BR) and oilseed rape only (OSR) data analyses. Note that GS had a differential effect on second-cohort fecundity (two filled circles): whereas GS had a positive effect on cross-species seed production, their presence did not alter oilseed rape fecundity. GS has been previously shown to deter seed predators (grey circle; de Jong *et al*. (2016).

In contrast to our cross-species analyses, differential investment in chemical defence had no noticeable effect on oilseed rape persistence (chemical PC1 in models 2 and 3: F_1,_75 = 0.47 and *χ*^2^_1_ = 0.30; *P* > 0.05; Table S3). Our results demonstrate that competition was a key determinant of population viability (soil PC1 in model 3: *χ*^2^_1_ = 36.69, *P* < 0.001; Table S3). Whereas cross-species recruitment was strongly affected by the availability of germination sites, vegetation turnover had little effect on oilseed rape recruitment (soil PC1 in models 1–3: F_1,72_ = 2.75, F1,73 = 2.65 and *χ*^2^_1_ = 2.85, all *P*> 0.05; Table S3). Instead, investment in EA/GS had a strong effect on seedling establishment, the extent of which varied across years (year x chemical PC1 in models 2 and 3: F_1,74_ = 36.05 and *χ*^2^_1_ = 31.75, all P < 0.001; Table S3) and localities (site x chemical PC1 in models 2 and 3: F_1,74_ = 6.12 and *χ*^2^_1_ = 6.36, all P < 0.05; Table S3). Patterns in plant survival were similar for both data sets: vegetation turnover had little effect on survival (soil PC1 in models 1–3: *P*> 0.05; Table S3), whereas the presence of EA/GS reduced plant survivorship (site x year x chemical PC1: F_1,67_ = 5.06 and *χ*^2^_1_ = 6.31, all P < 0.05; Table S3). In contrast to cross-species fecundity, oilseed rape plants produced more seeds in less densely vegetated patches (soil PC1 in models 2–3: *P*< 0.05 Table S3), independent of investment in EA/GS (models 2 and 3: P > 0.05; Table S3).

Results from the key factor analysis on the cross-species data set demonstrate that first-cohort *λ* was equally dependent on fecundity and recruitment in Swifterbant (*β*_*r*_=0.46^***^, *β*_*sa*_=0.07^NS^ and *β*_*F*_=0.47^***^), yet declines in *λ* were strongly associated with poor seedling establishment (*β*_*r*_=0.95^***^, *β*_*sa*_=-0.23^**^ and *β*_*F*_=0.28^*^). Demographic rates contributed equally, and positively, towards explaining variation in *λ* of the first cohort growing in Boskoop (*β*_*r*_=0.26^***^, *β*_*sa*_=0.35^***^ and *β*_*F*_=0.39^***^), whereas population decline in the second year was largely associated with reduced fecundity and adult survival (*β*_*r*_=0.17^***^, *β*_*sa*_=0.40^***^ and *β*_*F*_=0.43^***^). Together these findings suggest that selection for differential investment in EA/GS affects oilseed rape dynamics, by mediating trade-offs between recruitment and survival, which only became apparent during a second year of growth.

### Herbivory and predation

Herbivore damage was significantly lower for Swifterbant populations with greater investment in EA/GS, but only so in the second year of our study (chemical PC1 x year in GLM with quasipoisson error structure: F_1,53_ = 4.30, P = 0.04; Fig. 5A). The degree of pre-dispersal seed predation varied widely across Boskoop populations (Fig. 5B), independent of breeding history (1-way ANOVA on proportion of predated fruits: F_3,25_ = 1.83, P = 0.17) or investment in EA/GS (LME with chemical PC1 as fixed effect and random intercepts fitted for different levels of breeding history: *χ*^2^_4_= 3.30, P = 0.07).

**Figure 5.**
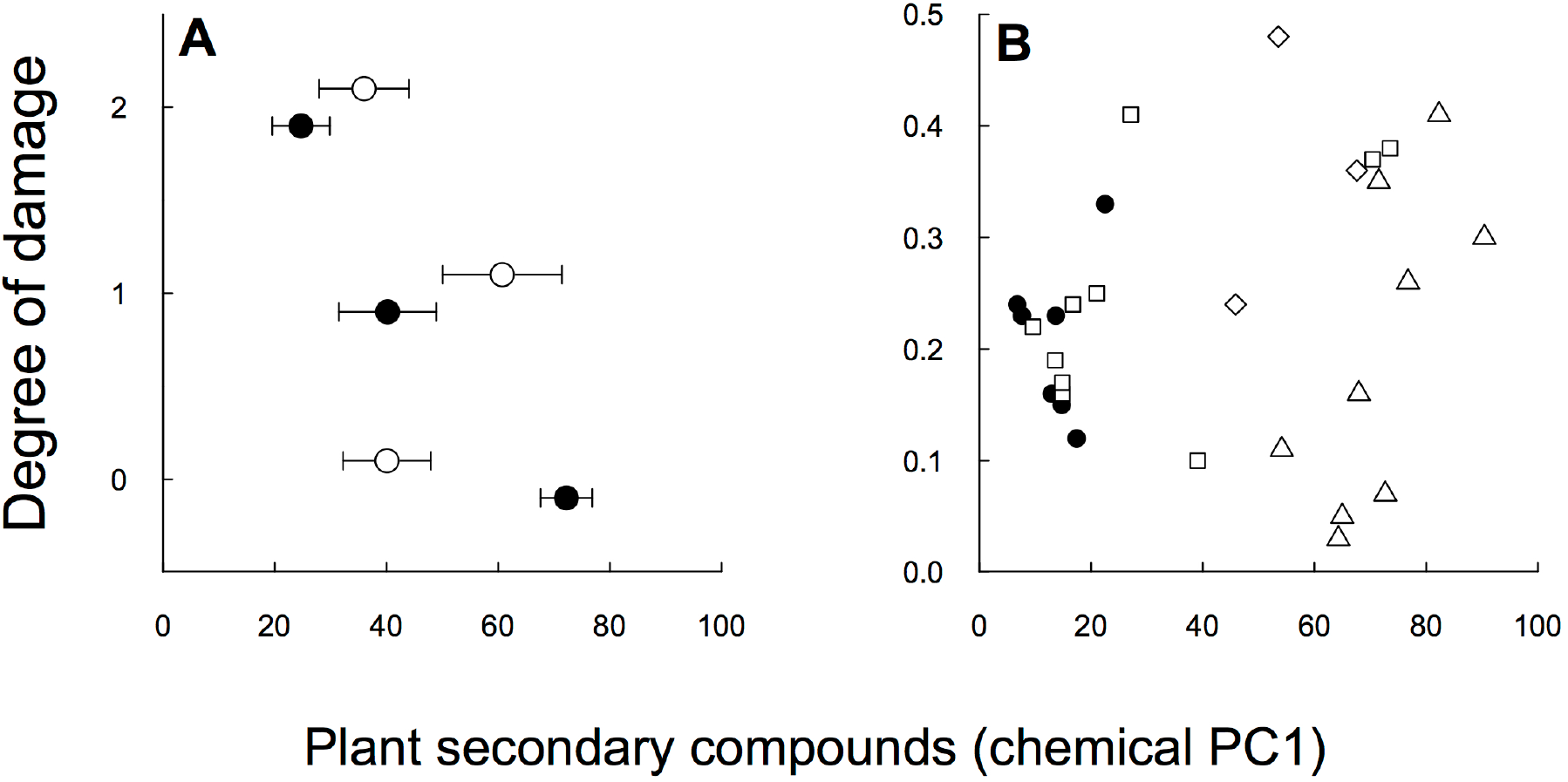
(A) Mean herbivore damage ± SE as a function of investment in secondary compounds (chemical PC1) during the first (open dots) and second (solid dots) year of growth in Swifterbant, and (B) Relationship between secondary compounds (chemical PC1) and the proportion of pre-dispersal seed predation in the first cohort of plants growing in Boskoop (dots = C, diamonds = CO, squares = BN and triangles = BR).

## Discussion

Glucosinolates have been linked to successful invasion of wild Brassicas (Muller 2009), yet the role of these compounds in determining oilseed rape ferality has been underappreciated. We aimed to unpack the risk of oilseed rape ferality into life stages, mediated by variation in GS and competition. Our results show that many populations perished during a second year of growth, in particular those of oilseed rape. Declines coincided with increased vegetation cover, yet were slower in populations harbouring more GS. Crucially, GS had opposing effects on underlying life cycle stages: GS enhanced seedling recruitment, yet benefits were negated following initial establishment as fewer high-GS seedlings reached the flowering stage. The net impact of GS on persistence was lost when studying oilseed rape only, but the underlying trade-off remained. Trade-offs only became apparent at the time when vital rates contributed differentially towards fitness: while recruitment was a key factor in relatively open habitats, survival contributed more towards fitness in more densely vegetated habitats. Because we did not measure environmental variables, such as humidity and nitrogen availability, we cannot rule out factors other than GS and competition important in driving ferality. Below we discuss potential mechanisms underlying the observed relationship between GS and various life stages.

Plants experience differential herbivory over the course of their life cycle (Boege & Marquis 2005). The consequences of herbivore attack, however, are often most serious for seedlings (Barton & Hanley 2013), which are primarily eaten by slugs and snails in temperate climates (Strauss et al. 2009). We hypothesized that variation in GS should alter the relative palatability of *Brassica* seedlings, with potentially cascading population-level consequences (Buschmann, Edwards & Dietz 2006). In line with previous experimental findings (Moshgani, Kolvoort & de Jong 2014), modern canola seedlings experienced particularly high levels of slug-induced mortality, particularly in the second year of our study. The fact that GS had little effect on the performance of first-cohort seedlings was likely due to temporary eradication of slugs and snails following tillage (E.H. personal observation). Recruitment not only encompasses successful germination and survival, but also avoidance of seed predators. While GS had little effect on pre-dispersal seed predation, variation in these compounds has been shown to alter seed attractiveness to granivores, with mice generally preferring low-GS seeds (de Jong et al. 2016). Crucially, without intensive monitoring, predation can be easily misconstrued as poor emergence resulting from competition (Strauss et al. 2009). Conversely, higher levels of GS decreased plant survivorship in the second year, suggesting that herbivore deterrence may come at a cost as a direct result of resource allocation constraints or, indirectly, by altering a plant’s competitive ability. Such costs of resistance have been detected in a wide range of plant species, including other Crucifers (Strauss et al. 2002).

While GS can protect plants from non-adapted generalists, specialist herbivores may use these compounds as oviposition or feeding cues, thereby exerting opposing selection on defence (Hopkins, van Dam & van Loon 2009). The selective value of chemical traits may therefore depend critically on the specific herbivore community in which plants grow, and may fluctuate over time, being favoured in years with relatively high generalist loads, and *vice versa*(e.g., Kliebenstein, Kroymann & Mitchell-Olds 2005; Lankau 2007; Gols et al. 2008; Chen et al. 2015). Moreover, a growing body of evidence suggests that GS play an additional role in plant-mutualist interactions (Kliebenstein, Kroymann & Mitchell-Olds 2005; Halkier & Gershenzon 2006): GS and their hydrolysis products can act as allelopathic compounds by disrupting mutualistic interactions between mycorrhizal fungi and seedlings, thereby indirectly suppressing growth of neighbouring plants (Stinson et al. 2006; Lankau & Strauss 2007; Muller 2009). Although we did not find any evidence for GS inhibiting growth of the residential community (data not shown), seedlings derived from high-GS lines did establish more successfully in relatively open plots. Hence, more experiments are needed to disentangle the relative roles of allelopathy *versus* herbivory in mediating seedling establishment.

Glucosinolate levels are known to vary both within and across *Brassica* populations (Hopkins, van Dam & van Loon 2009), yet little is known about chemical traits in feral oilseed rape. A subset of our feral populations displayed relatively high GS levels, which could have arisen as a result of founder effects (populations originating from high-GS cultivars), natural selection and/or hybridization (e.g., Charters, Robertson & Squire 1999; Pessel et al. 2001; Bond et al. 2004; Pivard et al. 2008; Elling, Neuffer & Bleeker 2009; Pascher et al. 2010; Schulze et al. 2014; Bailleul, Ollier & Lecomte 2016). The relative importance of these nonmutually exclusive processes in shaping trait variation remains largely unresolved (Elling, Neuffer & Bleeker 2009).

While effective herbivore deterrence depends on both chemical and morphological traits (e.g., Carmona, Lajeunesse & Johnson 2011; Chen, Gols & Benrey 2015; Chen et al. 2015), our findings strongly suggest that GS mediate interactions between *Brassica* plants and their generalist antagonists (e.g., Glen, Jones & Fieldsend 1990; Giamoustaris & Mithen 1995; Gols et al. 2008; Agrawal & Weber 2015). EA and GS concentrations were correlated in our study lines as a result of conventional breeding. However, EA is solely present in seeds and cotyledons; hence, variation in this fatty acid alone cannot explain differential herbivory in other life stages. Moreover, recent experimental work has demonstrated that variation in GS, rather than EA, has a major impact on predation of *Brassica* seeds by granivores (de Jong et al. 2016). Interestingly, plant volatiles can alter attractiveness of snails to modern canola seedlings, suggesting that olfactory cues may additionally influence plant-herbivore interactions (Shannon et al. 2016). A likely reason as to why these authors did not find a relationship between GS and snail feeding behaviour is the relatively low invariable GS levels present in their study lines, all falling below our detected range.

In line with previous work (Pekrun, Hewitt & Lutman 1998), few seeds developed secondary dormancy in the relatively undisturbed plots, which prevented the built-up of a seed bank. Several studies have demonstrated that oilseed rape seeds can become secondarily dormant following soil tillage (e.g., Gulden, Shirtliffe & Thomas 2003), or natural disturbances (Hooftman et al. 2015), the extent of which may vary across cultivars (Hails et al. 1997; Gruber, Pekrun & Claupein 2004). Hence, our measure of persistence likely underestimates longevity in more regularly disturbed habitats where recruitment from the seed bank has the potential to prolong local persistence (Hooftman et al. 2015). It would be worthwhile to quantify how differential investment in GS affects successful incorporation of seeds into a long-lived seed bank following disturbance, controlling for seed predation.

## Conclusions

We have revealed that GS may govern initial population establishment in semi-natural habitats by mediating trade-offs between recruitment and survival. As such, the relative importance of herbivory *versus* competition in determining oilseed rape ferality likely varies across the life cycle: low-GS lines should be more successful in habitats where survival is a key fitness component, whereas that of high-GS lines should be greater in habitats where recruitment contributes significantly towards fitness. Hence, the maintenance (persistence of relict plants) and/or evolutionary gain (hybridization and natural selection) of chemical traits may influence interactions between feral oilseed rape and their biotic environment. In order to increase our knowledge about the ecological role of GS in driving crop ferality, it is pertinent to investigate into more detail how these compounds may differentially affect life cycle transitions in a range of habitat types by altering ecological interactions with local herbivores (i.e., defence *versus* attraction) and/or competitors (i.e., indirect costs *versus* allelopathy).

## Acknowledgements

We thank André Kamp and Sonja Esch for help during data collection and the group of Nicole van Dam (Molecular Plant Physiology, Radboud University, The Netherlands) for carrying out glucosinolate assays and the lab of Christian Möllers (Division of Plant Breeding, University of Göttingen, Germany) for EA quantification. This work was supported by the Netherlands Organization for Scientific Research (grant number ERGO 383.06.112) and Natural Environment Research Council.

